# Comment on “TopHat2: accurate alignment of transcriptomes in the presence of insertions, deletions and gene fusions” by Kim et al.

**DOI:** 10.1101/000851

**Authors:** Alexander Dobin, Thomas R. Gingeras

## Abstract

In the recent paper [1] (thereafter referred to as “TopHat2paper”) the accuracy of TopHat2 was compared to other RNA-seq aligners. In this comment we re-examine most important analyses from the TopHat2paper and identify several deficiencies that significantly diminished performance of some of the aligners, including incorrect choice of mapping parameters, unfair comparison metrics, and unrealistic simulated data. Using STAR [2] as an exemplar, we demonstrate that correcting these deficiencies makes its accuracy equal or better than that of TopHat2. Furthermore, this exercise highlighted some serious issues with the TopHat2 algorithms, such as poor recall of alignments with a moderate (>3) number of mismatches, low sensitivity and high false discovery rate for splice junction detection, loss of precision for the realignment algorithm, and large number of false chimeric alignments.

## 1. Mapping real RNA-seq data

We utilized the experimental RNA-seq data ([3] GEO accession number: GSM818582) used in TopHat2paper, comprising ∼65M paired-end 2x101b reads.

### 1.1 Mapping parameters

For the TopHat2 runs, we used the same version (2.0.8) and parameters that were used in the TopHat2paper.

For all analyses in the TopHat2paper STAR was run with the following parameters: *–outFilterMismatchNmax 3 – outFilterMatchNmin 97 –outFilterScoreMin 90*. The presumed intention was to limit the number of mismatches to 3 per **each mate** of the paired-end read, since TopHat2 and other aligners in the TopHat2paper were allowed 3 mismatches per mate. However, – *outFilterMismatchNmax 3* parameter limits the total number of mismatches for the **whole paired-end read** rather than each mate. Hence STAR was only allowed to output reads with 3 mismatches per pair. For a consistent comparison, we ran STAR with – o*utFilterMismatchNmax 6,* i.e. allowing 6 mismatches per pair. The other two parameters used in the TopHat2paper, – *outFilterMatchNmin 97 –outFilterScoreMin 90,* have no effect on the mapping because they are overridden by the default values of –*outFilterScoreMinOverLread 0.66, –outFilterMatchNminOverLread 0.66.*

### 1.2 Edit distance

Following the TopHat2paper, we are using “edit distance” as the main metric of alignment quality. Edit distance of an alignment is defined as the total number of mismatched, inserted or deleted bases with respect to the reference genome. For multi-mapping reads, the alignment with the smallest edit distance is chosen. For TopHat2 alignments, we obtained the edit distance from the “NM” SAM attribute. For STAR, we calculated the edit distance by comparing the read sequences to the reference genome, counting mismatches for both mapped (“M” operation in CIGAR) and soft-clipped bases (“S” operation in CIGAR), and counting indels from “D” and “I” operations in CIGAR.

### 1.3 Genome and annotations

Following the Tophat2paper, we utilized ENSEMBL genome assembly and annotations. While reference genome in the TopHat2paper included only chromosomes 1-22,X,Y and mitochondrion genome (MT), we also incorporated non-chromosomal scaffolds (stored in the ENSEMBL file Homo_sapiens.GRCh37.66.dna.nonchromosomal.fa). We noted that the percentage of mapped reads for both STAR and TopHat2 are significantly higher in our runs than in the TopHat2paper, owing to a large number of reads that mapped to ribosomal RNA loci on the non-chromosomal scaffolds, which were not included in the TopHat2paper genomes.

### 1.4 Paired-end alignments

For paired-end reads, STAR standard mode of operation is to output only correctly (“concordantly”) paired alignments, while single-end and “chimeric” (discordant) alignments are filtered out. On the other hand, with the parameters used in TopHat2paper, TopHat2 was allowed to generate paired-end, single-end and chimeric alignments. In Section 2.2 we will show that TopHat2 produces a large number (∼10% of all reads) of false chimeric alignments even for a simplistic simulated dataset. It is possible that in real RNA-seq data, a few discordant pairs represent real fusion transcripts; however, the majority of discordant pairs are likely to be mis-mapping artifacts.

Therefore we believe the comparisons between aligners have to be performed only on concordantly mapped pairs. Several comparisons in the TopHat2paper (such as presented in Figures 2,3 / Tables S5,S6) do not take into account pair concordance and thus do not allow for a fair comparison.

**Figure 1.**
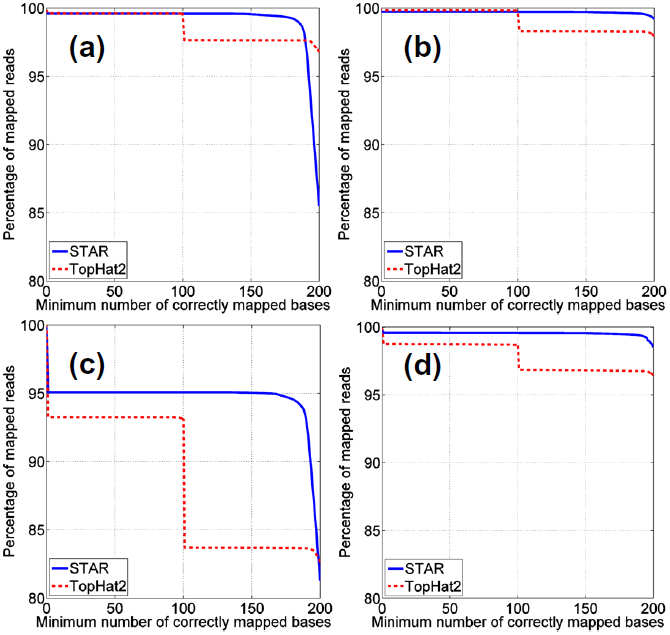
Percentage of reads (Y-axis) with the minimum number of correctly mapped bases per pair (X-axis) for simulated datasets. For each value *x* on the X-axis, the percentage of reads (normalized to the total number (20M) of simulated reads) mapped with the number of correct bases ≥ *x* is plotted. (a) Simulated data from the TopHat2paper: no realignment (Table 3, lines 1,2); (b) Simulated data from the TopHat2paper: STAR 2-step, TopHat2 realignment (Table 3, lines 3,4); (c) Simulated data with realistic gene expression: no realignment (Table 4, lines 1,2); (d) Simulated data with realistic gene expression, STAR 2-step, TopHat2 realignment (Table 4, lines 3,4).

In Table 1 we present the cumulative number of concordantly mapped pairs as a function of pair edit distance for the pairs (which is equal to the sum of edit distances of the mates). The same metric was used in TopHat2paper Figure S3 and Table S10. Columns 1 and 2 of Table 1 show the numbers of mapped reads for STAR and TopHat2 alignments without annotations, while columns 3 and 4 show the same numbers for annotations-aware alignments. In both cases STAR produces more alignments then TopHat2 for all edit distances, in stark contrast with the TopHat2paper results.

**Table 1.** Number of mapped pairs vs. pair edit distance.

| Column number |  | 1 | 2 | 3 | 4 | 5 | 6 |
| --- | --- | --- | --- | --- | --- | --- | --- |
| Method | Realignment | no (1-step) | no | no (1-step) | no | yes (2-step) | yes |
|  | Annotation | no | no | yes | yes | no | no |
|  | Aligner | STAR | TopHat2 | STAR | TopHat2 | STAR | TopHat2 |
| Maximum Edit Distance | 0 | 17,469,245 | 16,480,899 | 19,671,406 | 19,220,939 | 19,746,289 | 19,315,105 |
|  | 1 | 29,026,947 | 27,937,296 | 31,964,445 | 31,374,250 | 31,983,327 | 31,691,676 |
|  | 2 | 35,815,464 | 34,777,279 | 38,859,530 | 38,208,908 | 38,834,340 | 38,527,912 |
|  | 3 | 40,459,892 | 39,473,805 | 43,384,405 | 42,723,441 | 43,325,318 | 42,886,549 |

TopHat2 can be run in the “realignment” mode which substantially improves mapping sensitivity for annotations-lacking runs. As was discussed in the STAR paper [2], STAR can also be run in the 2-pass fashion, in which the junctions discovered in the 1^st^ mapping pass are utilized to increase sensitivity of the alignments in the 2^nd^ pass. While these advanced strategies improve performance of both aligners, the sensitivity of STAR 2-step mode is higher than that of TopHat2 realignment algorithm (columns 5 vs 6 of the Table 1). We note that this 2-pass scheme can be adapted by most RNA-seq aligners and is very beneficial for the situations where annotations are not available.

### 1.5 Alignments with moderate (>3) edit distances

The maximum edit distance was limited to 3 for all the analyses in the TopHat2paper. Realistic sequencing data, even of the highest quality, may contain reads with edit distances larger than 3, owing to poor quality of the read tails, poly-A tails, highly variable genomic loci, and RNA-editing. In addition, short splice overhangs often cannot be confidently positioned and have to be clipped off. As the read length grows with the advancements in sequencing technologies, the aligners will be required to deal with ever-increasing number of mismatches and indels. We ran STAR and TopHat2 with parameters allowing larger edit distances, and calculated the number of concordantly paired alignments as a function of pair edit distance (Table 2). Surprisingly, with the exception of the “realignment” mode, the numbers of TopHat2 alignments for edit distances from 0 to 3 dropped substantially (Table 2 columns 2,4) compared to the runs with more restrictive parameters (Section 1.4, Table 1 columns 2,4). Consequently, STAR’s mapping rate is substantially (>10%) higher than TopHat2’s in “no annotation” (columns 1 vs 2 of Table 2) and “annotation” (columns 3 vs 4 of Table 2) cases, and slightly higher for the “realignment” mode.

**Table 2.** Number of mapped pairs vs. pair edit distance for mapping that allowed for larger edit distances.

| Column number |  | 1 | 2 | 3 | 4 | 5 | 6 |
| --- | --- | --- | --- | --- | --- | --- | --- |
| Method | Realignment | no (1-step) | no | no (1-step) | no | yes (2-step) | yes |
|  | Annotation | no | no | yes | yes | no | no |
|  | Aligner | STAR | TopHat2 | STAR | TopHat2 | STAR | TopHat2 |
| Maximum Edit Distance | 0 | 17,469,245 | 14,606,065 | 19,671,406 | 17,871,997 | 19,746,289 | 19,286,070 |
|  | 1 | 29,026,946 | 24,884,770 | 31,964,444 | 29,049,298 | 31,983,326 | 31,662,265 |
|  | 2 | 35,815,460 | 31,045,091 | 38,859,526 | 35,274,349 | 38,834,334 | 38,487,634 |
|  | 3 | 40,459,752 | 35,285,388 | 43,384,270 | 39,363,178 | 43,325,245 | 42,823,798 |
|  | 4 | 43,940,148 | 38,470,105 | 46,662,374 | 42,364,584 | 46,569,290 | 45,833,817 |
|  | 5 | 46,681,608 | 40,954,184 | 49,141,568 | 44,664,377 | 49,015,510 | 48,025,776 |
|  | 6 | 48,896,786 | 42,985,446 | 51,067,990 | 46,472,697 | 50,915,421 | 49,689,282 |
|  | 7 | 50,683,823 | 44,659,572 | 52,602,413 | 47,936,908 | 52,427,060 | 50,991,882 |
|  | 8 | 52,148,832 | 46,103,633 | 53,850,418 | 49,141,737 | 53,656,184 | 52,052,668 |
|  | 9 | 53,343,317 | 47,375,766 | 54,895,487 | 50,141,346 | 54,684,000 | 52,922,650 |
|  | 10 | 54,337,233 | 48,476,866 | 55,770,256 | 50,988,912 | 55,542,821 | 53,657,365 |

## 2. Mapping simulated RNA-seq reads

### 2.1 Simulated dataset from the TopHat2paper

We mapped the simulated error-free paired-end 2x100b dataset utilized in the Table 2 of the TopHat2paper using the TopHat2 parameters from the TopHat2paper and default parameters for STAR.

In case of such error-free reads the choice of default or more restrictive parameters (as were used in the TopHat2paper) does not significantly affect STAR alignments. For each read we calculated the number of bases that were correctly mapped by each aligner. For the multi-mapping reads the alignment with the maximum number of correctly mapped bases was chosen.

Figure 1a shows the percentage of reads (Y-axis) with the minimum number of correctly mapped bases (X-axis). In agreement with observations in TopHat2paper, STAR finds completely (200b out of 200b) correct alignments for fewer reads than TopHat2 (85.5% vs 96.8%, lines 1 vs 2, column 1 of Table 3). As was pointed out in TopHat2paper, this is caused by STAR’s inability to align very short overhangs (∼10b or less) of spliced reads (referred to as “short-anchored” reads in the TopHat2paper). However, for the majority of these reads STAR aligns correctly the remainder of the sequence, and for reads with ≥190 correctly mapped bases (95% of the paired-end read sequence) STAR’s sensitivity reaches that of TopHat2. For almost all the reads in this simulated dataset, STAR maps correctly at least 150 bases, slightly outperforming TopHat2 (99.6% vs. 97.6%, lines 1 vs 2 column 2 of Table 3).

**Table 3.** Mapping statistics for the simulated RNA-seq dataset from TopHat2paper.

|  | Method |  | Correctly mapped<br>200<br>bases | >=150<br>bases<br>correctly<br>mapped | Unmapped | True positive<br>junctions |  | False positive<br>junctions |  |
| --- | --- | --- | --- | --- | --- | --- | --- | --- | --- |
|  |  |  |  |  |  | Number | Sensitivity | Number | FDR |
| Line<br>number | Realignment | Aligner | 1 | 2 | 4 | 5 | 6 | 7 | 8 |
| 1 | no (1-step) | STAR | 85.5% | 99.6% | 0.24% | 89,154 | 95.3% | 384 | 0.4% |
| 2 | no | TopHat2 | 96.8% | 97.6% | 0.14% | 84,178 | 90.0% | 1,268 | 1.5% |
| 3 | yes (2-step) | STAR | 99.2% | 99.7% | 0.26% | 89,234 | 95.4% | 270 | 0.3% |
| 4 | yes | TopHat2 | 97.9% | 98.3% | 0.14% | 84,838 | 90.7% | 3,511 | 4.0% |

Another important accuracy metric is the sensitivity and false discovery rate (FDR) of the splice junctions’ detection, presented in columns 5-8 of Table 3. Only uniquely mapped reads were considered, since including the multi-mapping reads improves sensitivity for both STAR and TopHat2 very slightly, while, at the same time, increases significantly the number of false positive junctions. Given that TopHat2 only detects junctions with GT/AG, GC/AG and AT/AC intron motifs, the junctions with all other intron motifs were excluded from this comparison. Even though STAR does not map very short spliced overhangs, it detects 4,976 more true junctions than TopHat2 (lines 1 vs 2 column 5 of Table 3) for a moderate gain in sensitivity (95.3% vs 90%, lines 1 vs 2 column 6 of Table 3). At the same time STAR demonstrates a much lower FDR than TopHat2 (0.4% vs 1.5% line 1 vs 2 column 8 of Table 3).

As we discussed above and in the STAR paper [2], in the absence of annotations, STAR will benefit from the 2-pass mode which improves the recovery of the short spliced overhangs by exploiting the junctions detected in the 1^st^ pass. This approach is compared to a similar TopHat2 “realignment” strategy in lines 3 vs 4 of the Table 3 and Figure 1b. The recovery of the completely correct alignments (200b) in the STAR-2-pass mode improves drastically to 99.2%, outperforming TopHat2-realignment rate of 97.9% (lines 3 vs 4, column 1 of Table 3). The junctions’ detection sensitivity of the TopHat2-realignment does not improve substantially; however, unexpectedly, the number of false positive junctions is severely increased (FDR=4%, column 8).

### 2.2 Simulated dataset with a realistic gene expression profile

Quoting from the TopHat2paper, “Both TopHat2 and MapSplice use a two-step algorithm, first detecting potential splice sites, and then using these sites to map reads. This two-step method may explain their superior performance at mapping reads with short anchors.” We note that such approaches (including STAR 2-pass) will work better if RNA transcripts are covered by a sufficient number of read, so that the “potential splice sites” can be confidently detected in the first step. However, real transcriptomes usually contain a large number of low expressed transcripts which do not satisfy this requirement.

To check whether the simulated datasets resembles the real RNA-seq data, we calculated the transcript abundances using the true read count per transcript for the TopHat2paper simulated dataset, and using Cufflinks 2.1.1 on TopHat2 alignments (run with annotations, no realignment, column 4 of Table 1) for the real RNA-seq data (same dataset as used in TopHat2 paper and the Section 1 of this Comment). Figure 2 shows the simulated and real distributions of FPKM values, defined as the number of paired-end fragments per kilobase of transcript length per million of mapped fragments. Compared with the real RNA expression profile, the simulated dataset is strongly enriched for high-abundance and depleted for low-abundance transcripts. This biased distribution is likely to favor “two-step” approaches such as TopHat2-realignment or STAR-2-pass.

**Figure 2.**
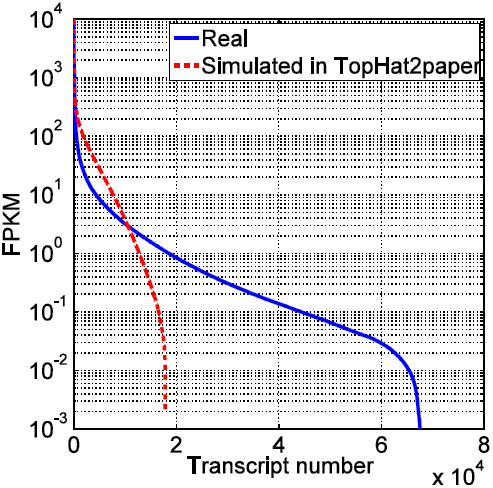
Transcript expression distribution for real and simulated RNA-seq data.

To assess the effect of the simulated transcript expression distribution bias on the mapping quality, we generated another simulated dataset with transcript abundances matching the real FPKM distribution. The number of simulated reads per transcript was proportional to real FPKM values with a total of 20 Million 2x100b paired-end reads. The 5’ ends of the reads were assumed to be distributed uniformly along the transcript sequences, while the 3’ ends were constrained by the 3’ ends of the transcripts, and required to produce a Gaussian distribution of insert sizes. Following the TopHat2paper, for simplicity of the comparison with Section 2.1, we did not introduce any genomic variations or sequencing errors in this dataset. STAR and TopHat2 were run on this more realistic simulated dataset and the results are presented in Figure 1c,d and Table 4. For the completely correct mapped reads, TopHat2-no-realignment mapping rate drops significantly to 82.6% (line 2 column 1 of Table 4) compared to the simulated dataset from TopHat2paper (line 2 column 1 of Table 3). STAR-1-pass achieves practically the same sensitivity as TopHat2 (81.3%, lines 1 column 1 of Table 4). At the same time STAR-1-pass achieves a significantly higher mapping rate than TopHat2-no-realignment for partially (≥150b out of 200b) correct alignments (95% vs 83.7%, lines 1 vs 2 column 2). STAR-1-pass also soundly outperforms TopHat2-no-relaignment in the splice junction detection sensitivity: 92.7% vs. 84.3% (lines 1,2 column 6) and junction FDR: 0.3% vs. 0.9%.

**Table 4.** Mapping statistics for the simulated RNA-seq dataset simulate with a realistic gene expression distribution.

|  | Method |  | Correctly mapped 200 bases | >=150 bases correctly mapped | Unmapped | True positive junctions |  | False positive junctions |  |
| --- | --- | --- | --- | --- | --- | --- | --- | --- | --- |
|  |  |  |  |  |  | Number | Sensitivity | Number | FDR |
| Line number | Realignment | Aligner | 1 | 2 | 4 | 5 | 6 | 7 | 8 |
| 1 | no (1-step) | STAR | 81.3% | 95.0% | 4.82% | 148,487 | 92.7% | 409 | 0.3% |
| 2 | no | TopHat2 | 82.6% | 83.7% | 6.70% | 135,006 | 84.3% | 1,228 | 0.9% |
| 3 | yes (2-step) | STAR | 98.5% | 99.5% | 0.40% | 148,789 | 92.9% | 453 | 0.3% |
| 4 | yes | TopHat2 | 96.4% | 96.8% | 1.22% | 136,416 | 85.2% | 3,132 | 2.2% |

The realignment (2-pass) strategy significantly improves performance of both aligners: the percentage of completely correctly mapped reads is increased to 98.5% for STAR-2-pass (line 3 of Table 4), which is slightly higher than 96.4% for TopHat2-realignment (line 4 of Table 4).

TopHat2 generates a large number of alignments for which only one mate is aligned correctly, which can be seen in Figure 1 as an abrupt drop in the percentage of reads with x=100 correctly mapped bases. Most of these alignments are false “chimeras” (alignments with mates mapping to different chromosomes or in wrong orientation). This effect is especially pronounced for TopHat2-no-realignment mapping of simulated data with realistic gene expression distribution (Figure 1c). In this case, even though the simulated data only contained perfect transcriptomic reads without mismatches or indels, TopHat2 produces ∼2 Million (10% of all reads) of false chimeras, revealing the danger of outputting unpaired alignments.

